# Parameter Identifiability of the Generalized Lotka-Volterra Model for Microbiome Studies

**DOI:** 10.1101/463372

**Authors:** Christopher H Remien, Mariah J Eckwright, Benjamin J Ridenhour

## Abstract

**Background:** Population dynamic models can be used in conjunction with time series of species abundances to infer interactions. Understanding microbial interactions is a prerequisite for numerous goals in microbiome research; predicting how populations change over time, determining how manipulations of microbiomes affect dynamics, and designing synthetic microbiomes to perform tasks are just a few examples. As such, there is great interest in adapting population dynamic theory for microbial systems. Despite the appeal, numerous hurdles exist. One hurdle is that the data commonly obtained from DNA sequencing yield estimates of relative abundances, while population dynamic models such as the generalized Lotka-Volterra model track absolute abundances or densities. It is not clear whether relative abundance data alone can be used to infer parameters of population dynamic models such as the Lotka-Volterra model.

**Results:** We used structural identifiability analyses to determine the extent to which time series of relative abundances can be used to parameterize the generalized Lotka-Volterra model. We found that only with absolute abundance data to accompany relative abundance estimates from sequencing can all parameters be uniquely identified. However, relative abundance data alone do contain information on relative interaction strengths, which is sufficient for many studies where the goal is to estimate key interactions and their effects on dynamics. Our results also indicate that the relative interaction rates that can be estimated using relative abundance data provide ample information to estimate relative changes of absolute abundance over time. Using synthetic data for which we know the underlying structure, we found our results to be robust to modest amounts of both process and measurement error.

**Conclusions:** Fitting the generalized Lotka-Volterra model to time-series sequencing data typically requires either assuming a constant population size or performing additional measurements to obtain absolute abundances. We have found that these assumptions are not strictly necessary because relative abundance data alone contain sufficient information to estimate relative rates of interaction, and thus to infer key drivers of microbial population dynamics.

## Background

There is considerable interest in applying population dynamic theory to microbial systems to test hypotheses relating to ecosystem stability, to determine the drivers of dynamics, and to predict how populations will change over time [e.g., to prevent illnesses such as ulcers; cf. 1–8]. While modern DNA sequencing technologies allow rapid and inexpensive characterization of microbial community composition and have uncovered enormous microbial diversity, relatively little is known regarding the interactions governing the population dynamics of constituent members of microbial communities. It is these interactions that determine which members of a microbial community will flourish, and understanding them is key to manipulating microbiomes to promote health, designing synthetic microbial communities to perform tasks, and inferring stability to assess risk.

Despite the expectation that population dynamic models should be applicable to microbial systems, barriers exist to the application of traditional modeling approaches to microbiomes. One such barrier results from the nature of sequence data as a proxy for species abundance. The raw data from high-throughput microbiome samples are a large number of sequence reads which are grouped by similarity, giving the number of reads belonging to a particular group. These groups have different meanings depending on the methods employed but, for our purposes, can be thought of as different (pseudo-)species. The number of reads for a group is then divided by the total number of sequence reads in the sample giving an estimate of relative abundance.

In contrast to the relative abundance estimates obtained from sequence reads, most population dynamic models, including the generalized Lotka-Volterra (gLV) model, describe absolute abundances or densities rather than sequence observation rates or relative abundances. Methods exist to convert the relative abundance data to absolute abundances by estimating absolute abundance from additional data (e.g., qPCR) [3, 9, 10]. However, such data are not typically collected in microbiome studies and, when collected, are quite error-prone themselves; thus we are often left with only estimates of relative abundances over time.

Numerous methods exist to estimate species’ interaction strengths in a gLV model from microbial time series data [3, 9–14]. The most common technique for estimating parameters is to utilize a discrete time version of the model, and estimate coefficients using linear regression. When formulated in this way, it has been recognized that the design matrix for the regression is singular [13] because relative abundance data alone do not contain sufficient information to estimate parameters. As such, methods typically rely on an assumption of constant population size or additional data on absolute abundance to complement sequence data so that absolute abundances or densities of each species can be estimated. While interaction strengths have been successfully estimated in microbial communities by fitting time-series of species’ densities to the gLV model, it is unknown to what extent relative abundances alone contain information on interaction strengths.

Using parameter identifiability analyses to systematically account for the compositional nature of the data, we determine the extent to which time-series of microbiome sequencing data contain information about parameters of the gLV model. We address the question as to whether relative abundance measurements alone, as obtained by sequencing techniques, can be used to estimate species interaction strengths in a gLV model. If relative abundance measurements cannot be used to estimate all species interaction strengths as has been previously suggested, what additional measurements would be needed to make this possible, and what parameters or combinations of parameters can be estimated using only relative abundance measurements? We then verify the structural identifiability results using synthetic microbial community time series data.

## Results

### Generalized Lotka-Volterra Model

Assuming large, well-mixed, closed populations with only two-way interactions between microbes, the change in density of microbes over time can be described by a system of differential equations where the dynamics of a focal microbe *N_i_* satisfy:

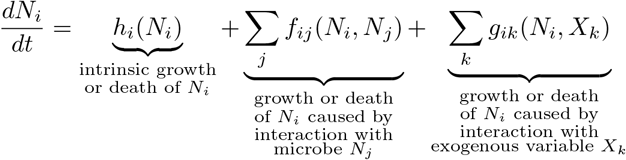

for *i* ∈ {1, 2, …*, n*}, where *n* is the number of species in the microbial community. The function *h_i_*(*N_i_*) is the rate of growth or death of *N_i_*, *f_ij_*(*N_i_, N_j_*) is a function describing the growth or death of *N_i_* caused by interaction with microbe *N_j_*, and *g_ik_*(*N_i_, X_k_*) is a function describing the growth or death of *N_i_* caused by interaction with exogenous variable *X_k_* (e.g., resource, toxin). Ignoring exogenous variables (*g_ik_* = 0) and specifying the functions *h_i_*(*N_i_*) = *r_i_N_i_* and *f_ij_*(*N_i_, N_j_*) = *β_i,j_N_i_N_j_* yields the classical generalized Lotka-Volterra model

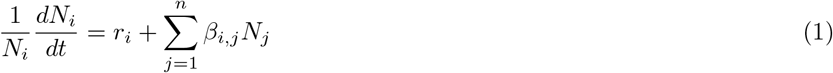

where the parameter *r_i_* is a positive growth rate and the interaction rate *β_i,j_* describes how microbe *j* affects the growth rate of microbe *i*. Typically, the parameters *β_i,i_* are constrained to be negative so that the carrying capacity *k_i_* = *−r_i_/β_i,i_* is positive.

### Structural Identifiability of gLV with Relative Abundance Data

A given model and observation state combination is said to be “structurally identifiable” if it is possible to uniquely estimate the parameters of the model assuming error-free measurements [15]. The goal of structural identifiability analyses is to identify model parameters that cannot be estimated from a given data type. Moreover, analysis of structural identifiability can reveal parameters or combinations of parameters that are uniquely identifiable, and can inform the re-parameterization of a model in terms of identifiable combinations of parameters. For the gLV model in equation (1), the structural identifiability problem can be set up as follows: given a noise-free time series of the relative abundance of each microbe (i.e., *N_i_/N* for all *i* where *N* = ∑*_i_ N_i_*), determine whether it is possible to estimate parameters *r_i_* and *β_i,j_* for all *i* and *j* of equation (1).

Results of structural identifiability analyses demonstrate that time series of relative abundance data do not contain enough information to uniquely estimate all parameters of the gLV model (equation (1)). Indeed, multiple sets of parameters can lead to identical relative abundance outputs (Figs. 1 and 2, Table 1). However, if additional information on absolute abundance can be obtained (e.g., through qPCR or optical density measurements) to complement relative abundance data, our results indicate that the parameters of a gLV model can be uniquely identified. Moreover, we demonstrate that such additional information need not be obtained at each time point; even one measurement of absolute abundance can, in theory, be used as the key piece of information to anchor the parameters giving identifiable estimates.

**Figure 1.**
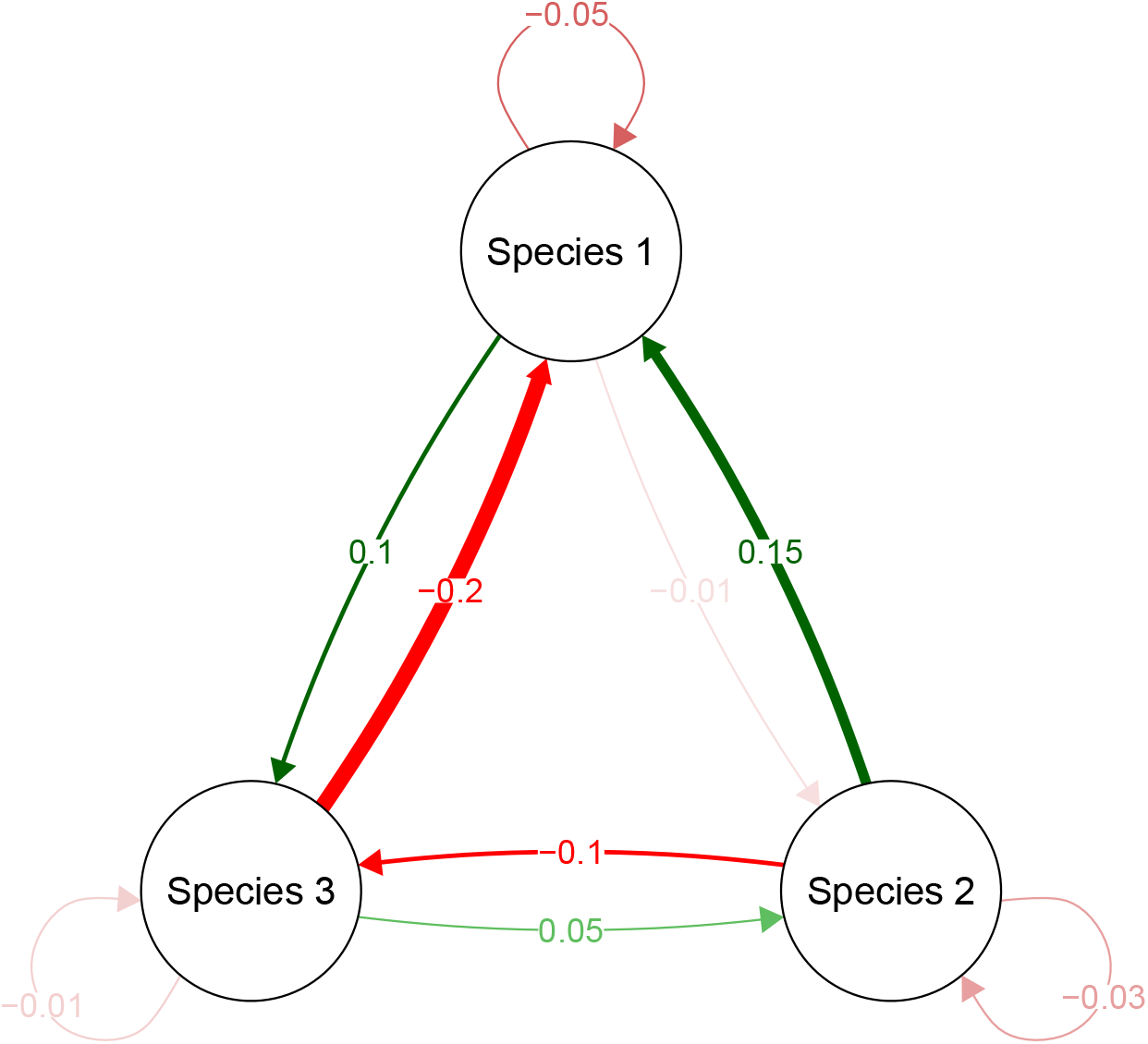
Interactions in the synthetic three-species community. The values in the arrows indicate parameters in the synthetic community.

**Figure 2.**
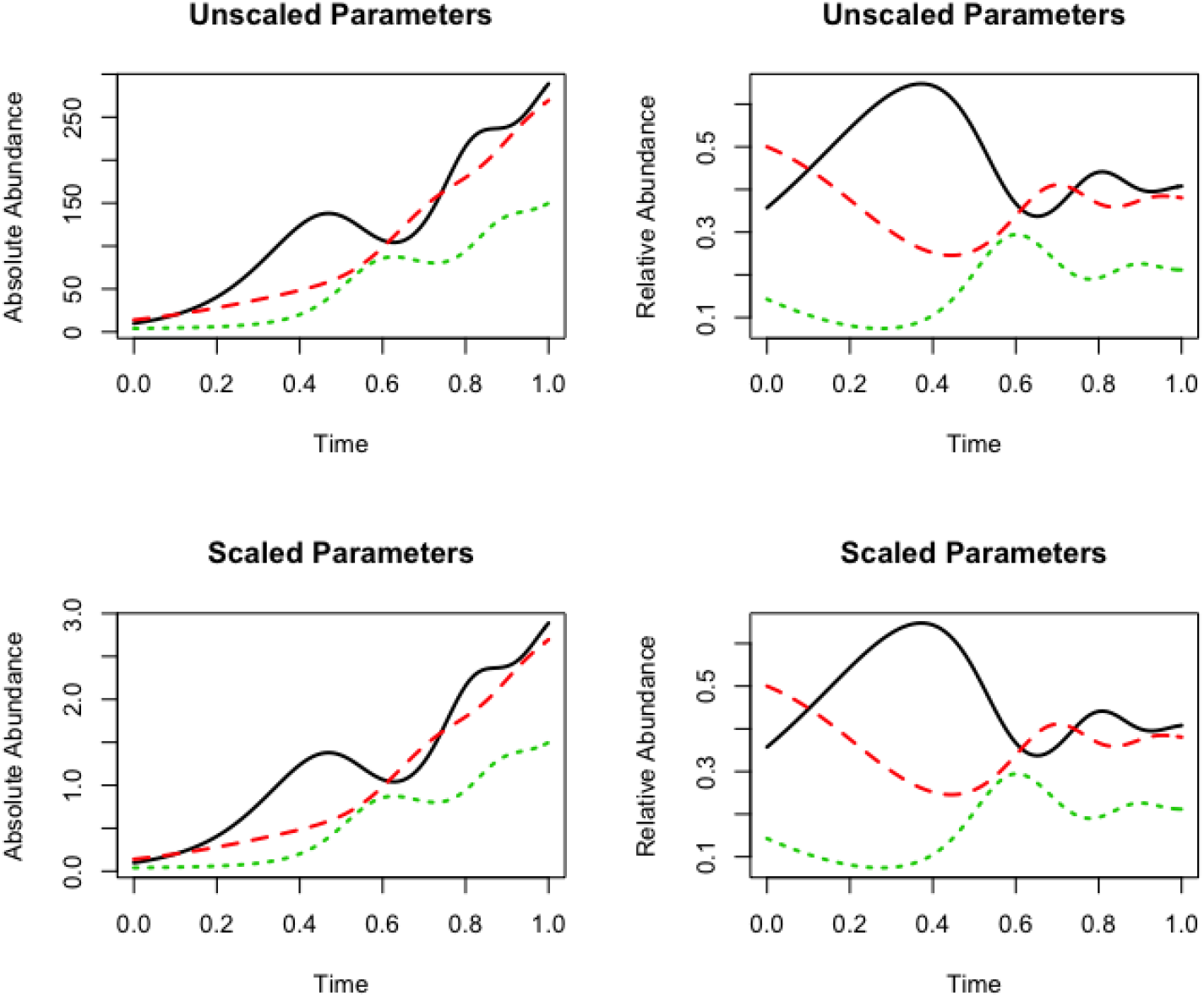
Multiple parameter sets lead to identical relative abundance dynamics. Comparison of two gLV systems with different parameters that yield different absolute abundances (left panels) but identical relative abundances (right panels). “Scaled” versus “unscaled” refers to the fact that the non-identifiable parameters can be scaled by a constant to produce infinitely many systems with identical dynamics in terms of the relative abundances. (The scaling constant chosen here was 100.) See Table 1 for parameter values.

**Table 1.**
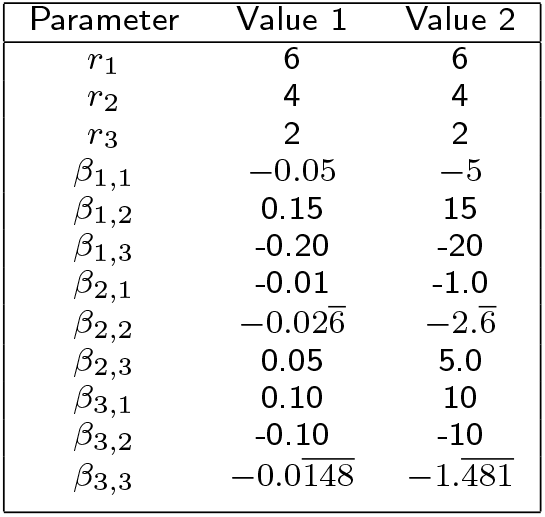
Parameter values used in numerical simulations

In most studies, additional information on absolute abundance is not available, and parameters of the gLV model must be estimated with only relative abundance data. While our results show that all parameters cannot be uniquely identified in this situation, we found that the relative interaction rates can still be obtained. The values of *β_i,j_* are identifiable up to a constant. Specifically, because the parameters *r_i_* are identifiable and *β_i,j_* are identifiable up to a constant, the relative topology of the interaction network corresponding to the interactions *β_i,j_* can still be estimated, even though the precise values of *β_i,j_* are not identifiable. Thus, given *relative* abundance data, *relative* interaction rates can be estimated. In many studies, this may be sufficient, as it would still allow for the identification of the key microbial interactions that provide services and drive dynamics.

### Application to Synthetic Community Data

To test whether modest observation and process error fundamentally affect the identifiability properties of the gLV model with relative abundance data, we created synthetic data of a three species community and sampled the likelihood surface of the parameter values given the synthetic data with noise (Fig. 3). We found that neither observation nor process error, nor both simultaneously, fundamentally affected the identifiability properties of the gLV model with relative abundance data (Fig. 4).

**Figure 3.**
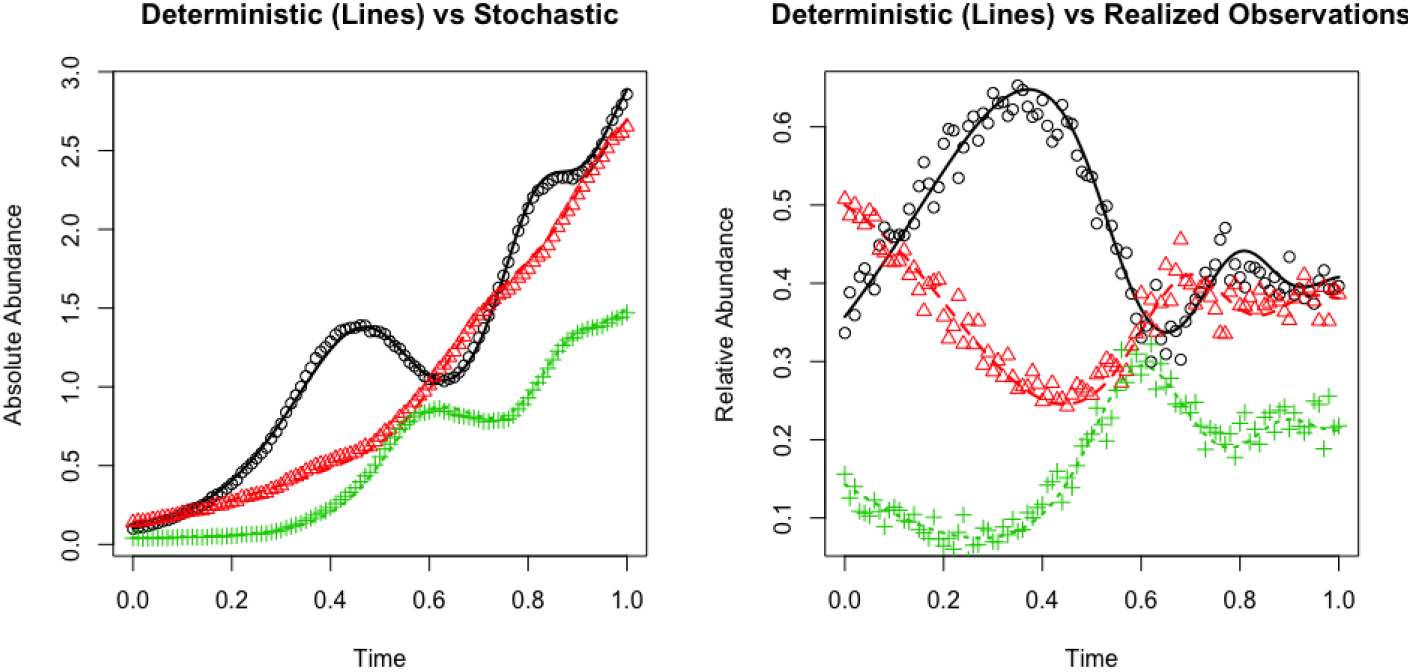
Dynamics with process and observation error. The example synthetic 3-species gLV system with process error and observation error incorporated. Process error was added using a Wiener process to simulate environmental noise. Observation error was added by sampling from a Dirichlet distribution with concentration parameters based on read-count number. See Methods for further details.

**Figure 4.**
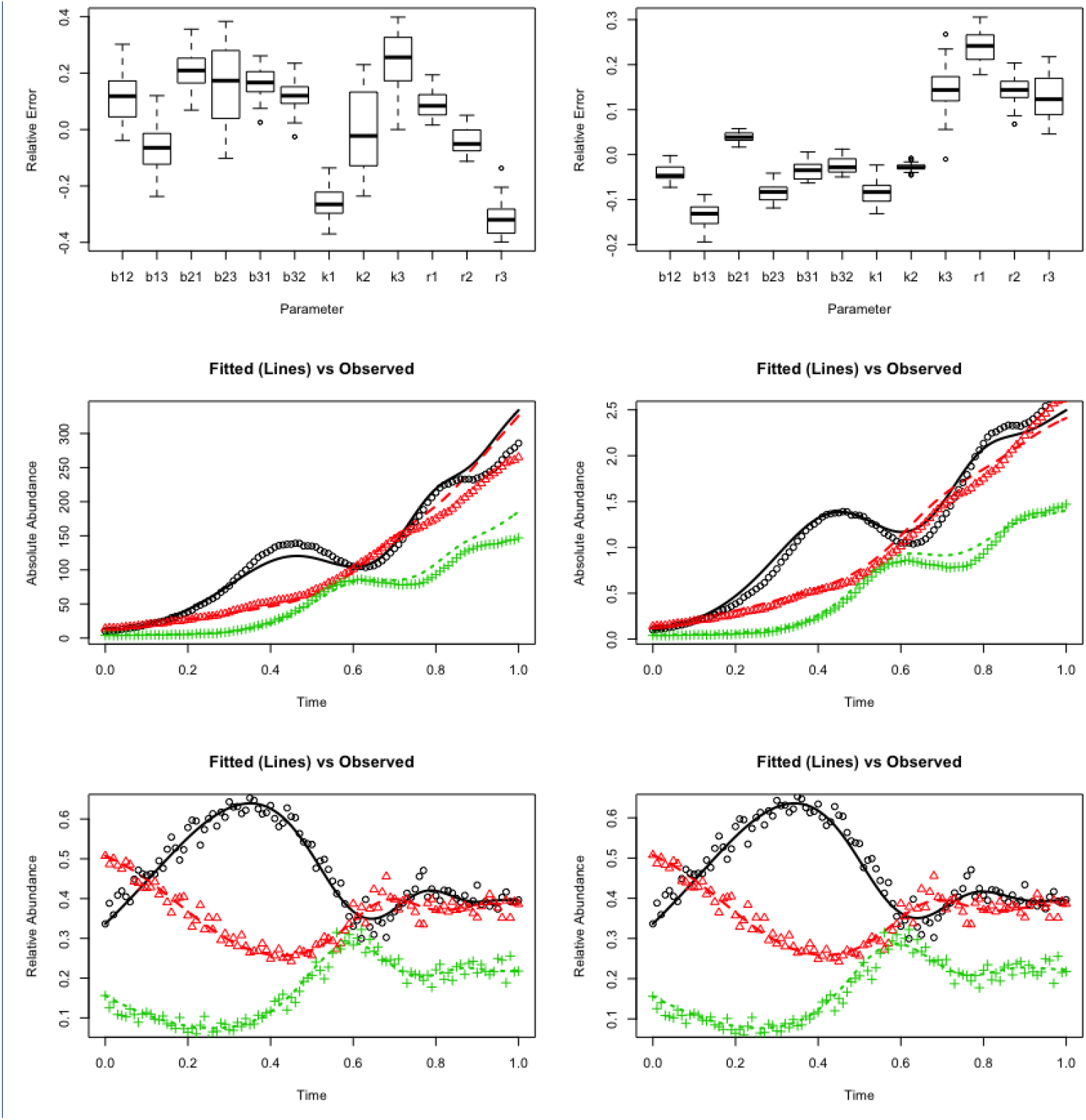
Fits of the synthetic 3-species gLV system with process and observation error. The left-hand column shows the results for the “Value 1” set of parameters (Table 1), while the right-hand column shows the results for the “Value 2” set of parameters. Fitting was done by starting the algorithm near the correct parameters set and giving it the correct total population size at *t* = 0 (*N*_0_ = 0.28 and *N*_0_ = 28, respectively). The fits for the relative abundance data are quite similar (last row), as are the extrapolated fits of the absolute abundance data (middle row). Relative error of the parameter estimates was defined as relative error = *x/x̂* − 1, where *x* is the estimated value and *x̂* is the true value. The relative error was bounded on [−0.4, 0.4] by the assumed priors.

Surprisingly, our results also indicate that the relative interaction rates—that can be estimated using relative abundance data—provide ample information to estimate relative changes of absolute abundance over time (Fig. 5). That is, total abundance relative to initial total abundance can be estimated from a time series of relative abundances of all microbes in conjunction with a gLV model. Of course, if the model is misspecified, or data sufficiently sparse, such estimation may not be possible for a given study system.

**Figure 5.**
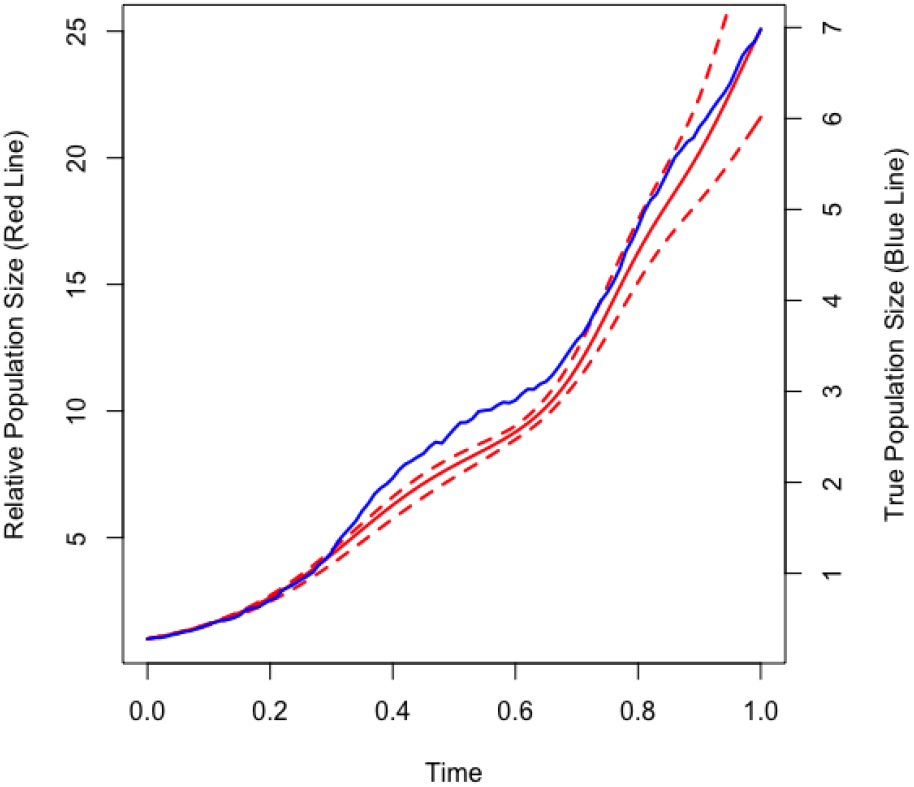
Reconstruction of relative population size. If a model is fit where it is assumed that *N*_0_ = 1, then information is recovered about the fold-change in the population’s size over time. The mean estimate (solid red line) does a fairly good job of matching the stochastic population trajectory (blue line). The dashed red lines show the range of population values produced by numerically solving the 3-species gLV system for all accepted parameter sets in the MCMC algorithm.

## Discussion

As a simple and general model that describes how interactions shape population dynamics of communities, the gLV is a natural candidate model for interpreting microbial time-series data [2, 4, 13, 16]. To date, fitting the gLV model to time series requires either 1) assuming a constant population size so that the gLV model can be fit directly to relative abundance data; or 2) performing additional measurements to obtain absolute abundances. We find that such assumptions or additional data are not strictly necessary. Using structural identifiability analysis and numerical simulations, we have shown that relative abundance data alone contain sufficient information to obtain relative rates of interaction. The ability to estimate the topology of an interaction network with only relative abundance time-series data would greatly expand the range of datasets available to interpret with dynamic models, as estimates of absolute abundances are typically unavailable.

Although our results demonstrate that much information can be gleaned from time-series relative abundance measurements, technical hurdles exist to efficiently estimate parameters of a gLV model with only relative abundance data. While multiple methods exist to fit gLV models to absolute abundance time-series data [e.g., 9, 10, 13], the most common technique is to substitute 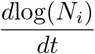 for 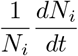 in equation (1), yielding

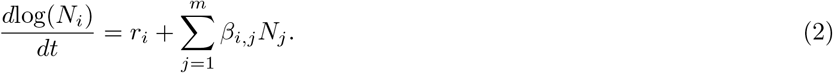

Because equations (2) are linear in *N_j_*, a discrete time version of the model can be fit to absolute abundance time-series data without the need to numerically solve the differential equations, typically with some form of regularization used to avoid overfitting. Unfortunately, there is no obvious analogous transformation of the relative abundance equations into a linear system, and thus to fit a gLV model to relative abundance data requires numerically solving the differential equations iteratively, which, while feasible for smaller systems, is a more computationally expensive proposition, particularly for large communities.

As with any statistical method, care must be taken when estimating parameters of the gLV model. Just because the relative interaction rates are structurally identifiable with relative abundance data does not mean that they are practically identifiable for all systems. Some parameters may not be practically identifiable due to the nature of the specific system. For example, if a species never gets near its carrying capacity, the carrying capacity may not be well estimated. Similarly, the number of data points is not the only determinant of the amount of information contained in a time series; time series in which populations exhibit large changes in population sizes due to, for example, perturbations, typically contain more information regarding interactions than time-series of species at steady state. The level and type of noise also dictates the ability to accurately estimate interaction parameters. For example, poor sequencing depth can increase measurement error and yield poor estimates of parameters.

Developing a better understanding of how microbes interact with each other and their environment is required for numerous goals in microbiome research, including detecting dysbioses, manipulating microbiomes to promote healthy function and preventing disease, and designing synthetic microbial communities for specific tasks. Such an understanding can be facilitated by interpreting data in conjunction with appropriate mathematical models. Some microbial communities may be better modeled with more complex formulations than the gLV model that incorporate additional factors, for example, higher order interactions, indirect resource-mediated interactions, time varying interactions, and various forms of stochasticity. The optimal level of detail to be included in such a model likely depends on numerous factors including the complexity of the microbial community, the level of understanding of the underlying dynamics, the structure of noise in the data, and the goals of the study.

## Conclusions

Regardless of the underlying model structure, population dynamic models must be adapted to utilize common forms of data, such as the relative abundance data obtained from high-throughput sequencing. We have found that when fit to a common population dynamic model, the generalized Lotka-Volterra model, a time series of relative abundance data contains information on relative interaction strengths. Moreover, relative interaction rates provide ample information to estimate relative changes of absolute abundance over time. Such findings provide critical information for designing temporal studies aimed at inferring microbial interaction networks, and greatly expand the number of studies amenable to such analysis. Specifically, we have shown that qPCR data—to convert relative abundance into absolute density—are not strictly necessary to obtain such networks. By appropriately connecting mechanistic models like the gLV with relative abundance data, we can potentially tease apart meaningful interactions governing microbial population dynamics.

## Methods

### Structural Identifiability of gLV with Relative Abundance Data

We wish to determine the upper bound on the information contained in a time series of relative abundance data in relation to the gLV model defined in equations (1). As previously mentioned, after grouping by similarity, sequencing data give estimates of the relative abundance of each group, whereas equation (1) tracks absolute abundances or densities. We will use structural identifiability analyses to determine the extent to which relative abundance data can be used to infer population dynamic parameters.

A variety of methods exist to determine structural identifiability of ODE models [17]. We will utilize a differential algebraic approach which is relevant for rational-function ODE models such as equation (1). To apply this method on the model described in equation (1), the idea is to first algebraically manipulate the system of differential equations into an equivalent system written only in terms of *observable* state variables (i.e., the measured data) and their derivatives. The resulting system can be regarded as a system of differential algebraic equations (DAEs) with polynomial coefficients which, after dividing by the coefficient of the highest ranking polynomial to make the resulting system monic, leads to an input-ouput relation that has identifiable coefficients [15, 18]. While this method is theoretically valid for a microbial community of arbitrary size, the algebra becomes cumbersome for even relatively small communities. Nevertheless, analyses of communities of small size uncovers clearly recognizable patterns that appear to be broadly applicable to communities of arbitrary size.

We begin by analyzing the structural identifiability of the two species gLV model

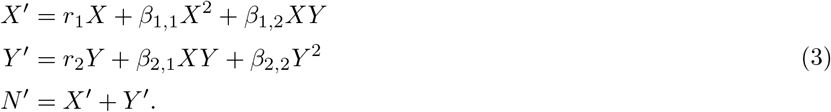

In equations (3), *X* represents the absolute abundance of the first species, *Y* the absolute abundance of the second species, *N* = *X* + *Y* the total abundance of microbes in the community, and all derivatives are with respect to time. Note that *N, X,* and *Y* are all time-dependent (e.g., *N* (*t*)), but we are suppressing the time notation for brevity. Ideally, we would like to use measurements of relative abundances to estimate parameters of (3). To check identifiability, we rewrite equations (3) in terms of the measurable quantities (relative abundances). Let *x* = *X/N* and *y* = *Y/N* be relative abundance of microbes *X* and *Y*, respectively. Differentiating *x* with respect to *t* yields

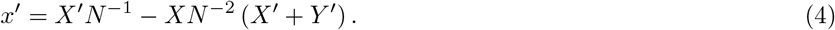

Utilizing *X* = *N_x_* and *Y* = *N* (1 *− x*) and equation (4), equations (3) can be rewritten in terms of relative abundances and *N* as

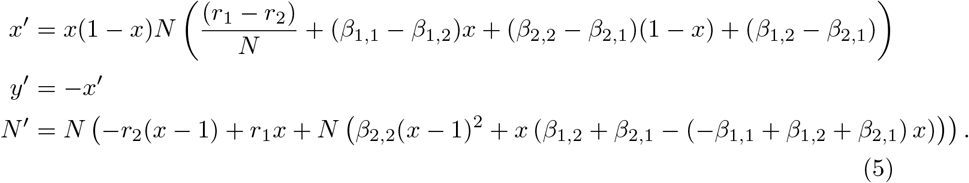

Solving the first equation in (5) for *N* yields

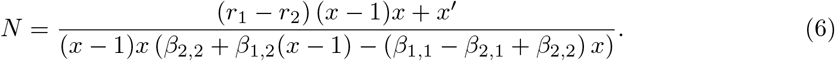

Substituting (6) and its derivative into the *N*′ equation in (5) and collecting terms yields

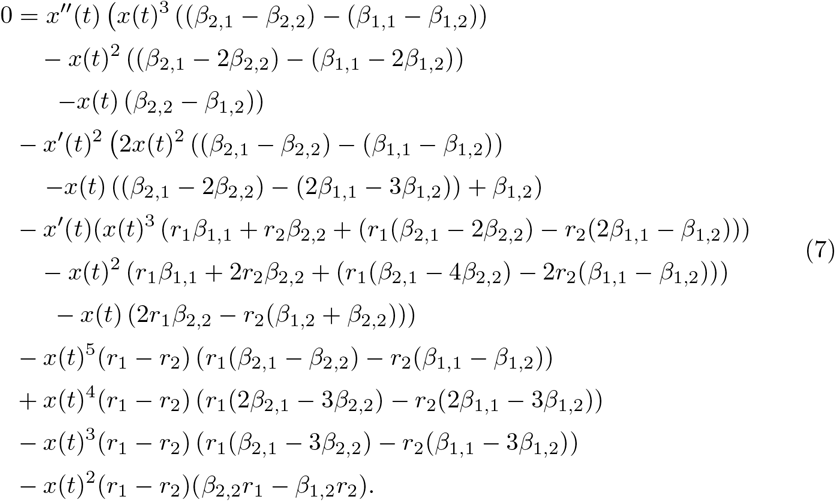

Dividing equation (7) by the leading order coefficient ((*β*_2,1_ *− β*_2,2_) *−* (*β*_1,1_ *− β*_1,2_)) produces an input-output DAE strictly in terms of *x′′*, *x′*, and *x* whose coefficients are identifiable. Finally, we check whether coefficients of the DAE have a unique solution by considering an alternative set of parameters (*a*_1_*, a*_2_*, a*_3_*, a*_4_*, a*_5_*, a*_6_) that produces the same output. Doing so gives the following result: *r*_1_ = *a*_1_, *r*_2_ = *a*_2,_ 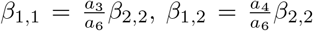, and 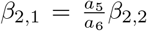. Thus, *r*_1_ and *r*_2_ are identifiable using relative abundance data, while *β_i,j_* are not identifiable; however, *β_i,j_* are identifiable up to a constant.

We performed similar analyses for three, four, and five species gLV models and found similar identifiability results (Supplemental Mathematica Notebook). Parameters *r_i_* are identifiable, while *β_i,j_* are only identifiable up to a constant.

### Application to Synthetic Community Data

To verify the structural identifiability results of the previous section and to test how error affects the identifiability results, we created two synthetic communities that, according to the structural identifiability analysis in the previous section, should yield identical relative abundance time series (Table 1, Fig. 1). We numerically solved equations (3) with parameters in Table 1 using the function ode with the default lsoda method in the deSolve R package.

To determine how error affects the identifiability results, we created more realistic synthetic data with two sources of error: process error and measurement error. Process error was added by using stochastic differential equations. Specifically, we used a Wiener process (Brownian motion) to model environmental noise to the system [19]:

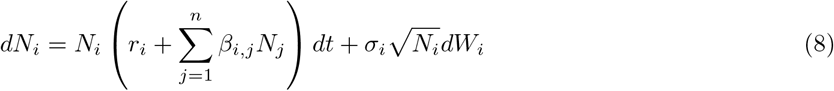

where σ_*i*_ scales the variance of the Wiener process (*dW_i_*), which is 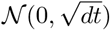- distributed random noise. After addition of the process error, we subsequently added measurement error to the stochastically-modeled relative abundances. To do this, we drew random proportions from a Dirichlet distribution having concentration parameters *α_i_* = *V Ñ_i_/*∑*_i_ Ñ_i_*, where the tilde indicates the population sizes simulated by Eq. (8) and V scales the error magnitude (V is roughly equivalent to the amplicon read count). Simulation of data with process and observation error was performed using the pomp function in the POMP R package [20].

Once we created our synthetic data, we then used POMP’s particle Markov chain–Monte Carlo (pMCMC) algorithm [cf. 21] to see if the correct parameters could be inferred for several scenarios: the two scenarios presented in Table 1 and a third scenario where the initial population size *N*_0_ = 1. The third scenario was performed to demonstrate that *relative* changes in population size can be inferred from relative abundance data (i.e., *N* (*t*) *≈ N*_0_*N̂*(*t*) where the hat denotes the estimated population size). Because we are strictly interested in whether the likelihood surface has a maximum at the correct parameter values, we began the algorithm near the correct parameter set to verify convergence of the fitting algorithm. There may be parameters that fit the realization with both types of error better than the parameters that created the realization; however, the fitted parameters should be similar to the parameters used to create the simulation.

The fitting was performed as follows. We assumed that every sample had 500 amplicon read counts that were then divided with respect to the relative proportions of each species; accordingly, we specified that read counts (measurements) were multinomially distributed. The time-step (*dt*) for the random process error was 0.001. A prior uniform distribution was placed on each parameter, *ρ*, such that the likelihood surface was defined on 𝒰(*ρ −*0.4*|ρ|,ρ*+0.4*|ρ|*). The MCMC algorithm was first run for 2000 iterations with 200 particles used for filtering at each iteration. During these iterations proposals were drawn using a multivariate-normal, adaptive, random walk where the covariance matrix of MVN was defined as a diagonal matrix with non-zero elements corresponding to 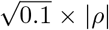; after 100 iterations, a scaled empirical covariance matrix based on the accepted proposal was used. Once the first 2000 iterations finished, the MCMC sampler was restarted and run for another 2000 iterations using the empirically determined covariance matrix from the previous 2000 iterations. After sampling was completed, the final 2000 iterations were thinned by keeping only every 50^th^ sample, resulting in 40 proposals from which to estimate the summary statistics for each parameter.

## Declarations

### A. Ethics approval and consent to participate

Not applicable

### B. Consent for publication

Not applicable

### C. Availability of data and material

Data sharing not applicable to this article as no datasets were generated or analyzed during the current study. Code is available as additional files.

### E. Author’s contributions

CHR and BJR designed the research. CHR, MJE, and BJR performed mathematical and computational analyses. CHR and BJR drafted the manuscript with input from MJE.

## F. Acknowledgements

We thank the CMCI Microbiome working group for useful discussion.

## Additional Files

Additional file 1 — Mathematica Notebook

Mathematica notebook for parameter identifiability of 2-5 species generalized Lotka-Volterra models with relative abundance data.

Additional file 2 — R Markdown File

R code for the application to synthetic community data.

